# Predictive Coding algorithms induce brain-like responses in Artificial Neural Networks

**DOI:** 10.1101/2025.01.16.633317

**Authors:** Dirk Gütlin, Ryszard Auksztulewicz

## Abstract

This study explores whether predictive coding (PC) inspired Deep Neural Networks can serve as biologically plausible neural network models of the brain. We compared two PC-inspired training objectives, a predictive and a contrastive approach, to a supervised baseline in a simple Recurrent Neural Network (RNN) architecture. We evaluated the models on key signatures of PC, including mismatch responses, formation of priors, and learning of semantic information. Our results show that the PC-inspired models, especially a locally trained predictive model, exhibited these PC-like behaviors better than a Supervised or an Untrained RNN. Further, we found that activity regularization evokes mismatch response-like effects across all models, suggesting it may serve as a proxy for the energy-saving principles of PC. Finally, we find that Gain Control (an important mechanism in the PC framework) can be implemented using weight regularization. Overall, our findings indicate that PC-inspired models are able to capture important computational principles of predictive processing in the brain, and can serve as a promising foundation for building biologically plausible artificial neural networks. This work contributes to our understanding of the relationship between artificial and biological neural networks, and highlights the potential of PC-inspired algorithms for advancing brain modelling as well as brain-inspired machine learning.

## 1. Introduction

The relationship between biological and artificial neural networks is a topic of growing importance, as insights from each field can inform and advance the other. While deep neural networks (DNNs) have emerged as a powerful model for investigating brain processes [1,2], their components and learning mechanisms are not directly aligned with our understanding of biological neural networks [3–8].

### 1.1. What exactly are the limitations of current neural network models as brain models?

DNN models are essentially machine learning constructs, typically optimized for specific tasks rather than biological plausibility. Consequently, these models rely on learning mechanisms and architectural choices that are only loosely aligned with our knowledge of biological neural networks [3–8]. A key example is supervised backpropagation, which is ubiquitous in deep learning. ‘Supervised’ means networks are provided with an externally provided ground truth, which is implausible for biological neural networks [9]. ‘Backpropagation’ refers to the updating rule used for DNN which updates model parameters hierarchically from target to input, calculating partial derivatives at each layer and propagating them back [10]. In biological networks, such a mechanism becomes increasingly unlikely the more layers are involved [3,11–15]. Despite being powerful functional models, current DNN models (especially when trained with supervised backpropagation), are not plausible models of biological neural networks.

While past neuroconnectionist research has largely focused on architecture [16–23], more recent work has shown that training objectives play a comparably large role [24]. For example, in visual object recognition neuroscience, models have traditionally used supervised objectives and backpropagation-based optimizers [16–23]. However, unsupervised objectives like SimCLR [25] have emerged as more biologically plausible alternatives that can yield equally effective or better representations [24,26]. Nevertheless, these unsupervised rules are still often based on information-theoretical ideas rather than biological principles. A promising avenue of research is to look to neuroscience for inspiration on biologically plausible learning mechanisms for DNNs. Several such approaches have been proposed.

1. Hebbian learning rules, inspired by synaptic plasticity, are intuitively appealing but often too crude to learn complex relations [27–29].
2. Reinforcement learning aims to capture reward-based learning, but current models are not as effective as classical deep learning optimization [30–33].
3. Bayesian learning rules provide a convenient connection between probability/information theory and neuroscience. Bayesian learning [34,35] in general treats the brain as a ‘prediction machine’, by actively predicting future states and trying to optimize these predictions. The family of Bayesian learning rules includes more specified theories of neural learning, such as the free energy principle [36,37] and predictive coding [38,39]. These theoretical frameworks are more directly inspired by empirical results from neurophysiology and provide sufficient detail and operationalization to actually implement it in a computer. Due to their biological plausibility and clear cut formalization, they provide a well-suited candidate framework for building biologically plausible artificial neural networks.

In this research article, we will therefore focus on predictive coding models as the most suitable class of biologically plausible DNN models of the brain.

### 1.2. What is predictive coding?

Predictive coding (PC) posits that the brain is fundamentally engaged in generating predictions about incoming sensory inputs and updating internal models based on the mismatch between predictions and observations. The primary function of PC is to approximate the prediction (prior) as closely as possible to the actual future stimulus (posterior), enabling timely adaptation to new circumstances. Further, as an energy-minimizing procedure, PC proposes that only the residual, unpredicted information should be propagated to higher levels, as the expected information can be suppressed. The main computational operation is a subtraction between the prediction and the incoming stimulus (although more recent formulations also emphasize multiplicative operations such as the precision of prediction errors, linked to biological concepts such as neural gain control [40,41]). This PC framework has gained significant traction in neuroscience, as it provides a principled account for phenomena like hierarchical representations and error signals driving learning and adaptation [38–43].

The most popular PC-inspired algorithm is PredNet by Lotter et al. [44]. PredNet combines a convolutional neural network (CNN) with a long short-term memory (LSTM) recurrent layer and an autoregressive/predictive target, resulting in good performance on video prediction tasks. Recently, an alternative model - namely the Forward-Forward algorithm - has been proposed by Hinton [45]. It integrates ideas from Boltzmann machines, generative adversarial networks, and contrastive learning to form a training setting that can resemble the PC mechanism under the right conditions. Additionally, a range of other algorithms have been proposed, including work such as Millidge et al. [11,46] or Whittington & Bogacz [47]. These approaches often consist of custom architectures and custom updating rules, with errors explicitly calculated. Despite the wealth of different existing predictive-coding inspired DNN, only few of these models have ever been used as a model for the biological brain so far [44,48]. While using DNN models as models of brain function is currently growing in popularity, this field of research usually focuses on more functionally performant, but less biologically plausible models. Accordingly, it is mostly unknown to what extent these biologically plausible DNN algorithms realistically approximate neural responses.

### 1.3. What is the main goal of this work?

This work aims to investigate whether PC inspired algorithms match phenomena or brain data better than classical supervised backpropagation in a simple perceptive setting. This would situate PC inspired algorithms as a promising alternative to other less biologically plausible approaches. To reach this goal, we compare PC inspired optimization approaches applied to one common simple Recurrent Neural Network (RNN) architecture (consisting of a feedforward input kernel and one square recurrent kernel). Since here we aim at comparing general PC-inspired algorithms and updating rules in a well-controlled setting, we focus on two candidates: A Predictive implementation, inspired by PredNet [49] and a Contrastive implementation, based on Forward-Forward [45]. We train these methods on a simple visual perception task, as a sufficiently complex and intuitively understandable task to demonstrate the effects of PC. As previous literature suggests that applying activity regularization to a network can by itself introduce PC-like effects in a network [50], we introduce an additional condition, where each of these models is trained with activity regularization applied next to the main objective. We then test whether these PC-inspired neural network models exhibit key neural signatures of predictive processing, in comparison to standard supervised and untrained neural networks. Specifically, we define three necessary landmarks which a PC network should exhibit:

1. Mismatch responses (MMR), which reflect the network’s ability to detect and respond to unexpected or deviant stimuli, and is core to the concept of saving energy in PC literature [51–53].
2. Prior expectations (i.e. actively maintaining predictions about upcoming states), which are a necessary precondition for a PC system [54–60].
3. Learning of semantic information (i.e. the association of a stimulus to an abstract encoded concept over the course of training) without explicit supervision. This is a necessary attribute of any learning system.

We use these landmarks as dependent variables, quantifying to what extent the mentioned phenomena emerge in different model types (see Figure 1). Furthermore, since gain control is considered a crucial component of PC and enables flexible context-sensitive weighting of prediction errors [40,41], we perform an additional analysis to investigate whether ANN weight regularization can be used to simulate gain control.

**Figure 1:**
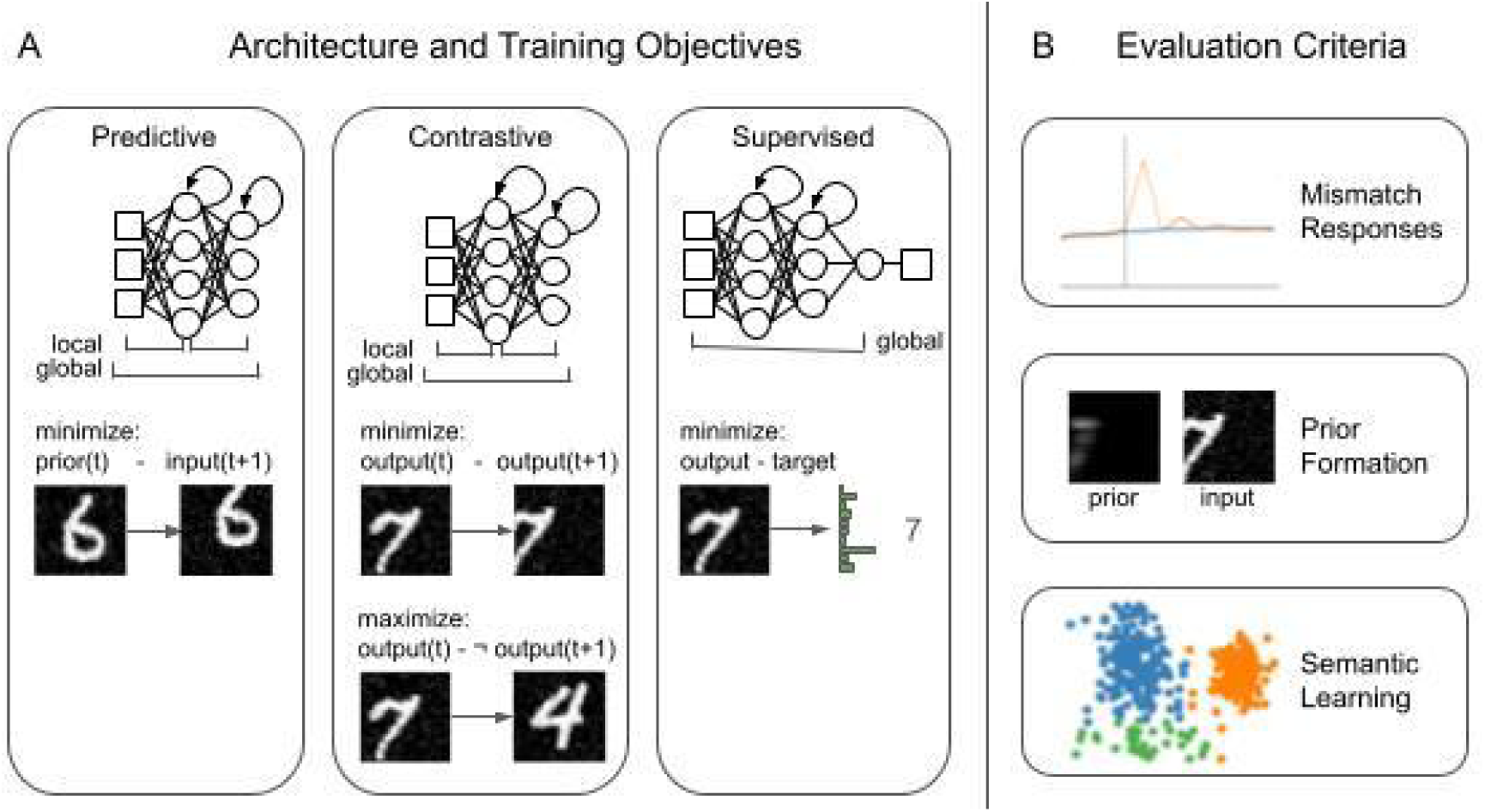
Illustration of the different training objectives (main experimental conditions) and evaluation criteria (dependent variables). A. Network Conditions: Depiction of each training condition (Predictive, Contrastive, Supervised), illustrating the simple Recurrent Neural Network Architecture (RNN) used as well as the optimization objective of each network. For the predictive algorithm, the supervised objective can only be applied using “global” gradient propagation (backpropagation) through the entire network. However, the Predictive and the Contrastive objective can be applied on a “local” scale, with each layer being trained on the inputs and outputs of the current layer. B. Evaluation criteria: Illustration of the three dependent variables to assess phenomenological hallmarks of PC in the networks.

## 2. Results

To investigate which neural network model aligns most closely with PC, we compared PC-inspired training objectives (mentioned in Introduction 1.3.) to a Supervised and Untrained baseline. We created several different training conditions:

1. Predictive global, where a predictive loss is applied after the final network output.
2. Predictive local, where a predictive loss is applied after each layer.
3. Contrastive global, where a contrastive loss is applied after the final network output.
4. Contrastive local, where a contrastive loss is applied after each layer.
5. Supervised, a control condition where a supervised loss is applied after the final layer.
6. Untrained, a control condition in which the network is not trained at all.

All conditions were trained on identical series of images (for details see Methods 4.2.) in an identical simple RNN architecture (see Figure 1). While the Supervised condition was trained to correctly identify the number presented in each image of the series, the PC-inspired conditions were trained according to unsupervised PC objectives (for details see Methods 4.3.1.). We then compared all six training conditions in respect to the hallmarks of predictive coding defined in Introduction 1.3.: (1) generation of mismatch responses, (2) formation of prior expectations, and (3) learning of class-specific semantic information. In the following section, we review each evaluation criterion and explain which model condition shows the corresponding hallmarks of PC. Since activity regularization may also evoke PC-like dynamics in networks, we first present the results for networks trained without activity regularization, and then for those trained with additional activity regularization.

### 2.1. Mismatch responses

In PC, the occurrence of MMR is directly tied to deviations from what was expected, resulting in elevated activity [61]. We expect similar MMR patterns in PC-inspired networks. To assess the models’ ability to detect and respond to unexpected or deviant stimuli, we evaluated their MMR (measured by deviations in the evoked neural response) to a series of matching and mismatching input sequences (see Figure 2). The matching series condition consisted of a sequence of MNIST images with random augmentations, while the mismatching series condition included random augmentations and random switches in the semantic content of the images (e.g., a 3 suddenly changing to a 7). We ensured that the match and mismatch condition did not show any base level differences that might result in a difference in mean-squared activation from the network’s layers (for details see Methods 4.4.1.). We then measured the mean square layer activations of the networks over time, similar to evoked responses. Our metric of interest was whether the match and the mismatch condition produced significant deviations in the model output for the sample immediately following the stimulus switch.

**Figure 2:**
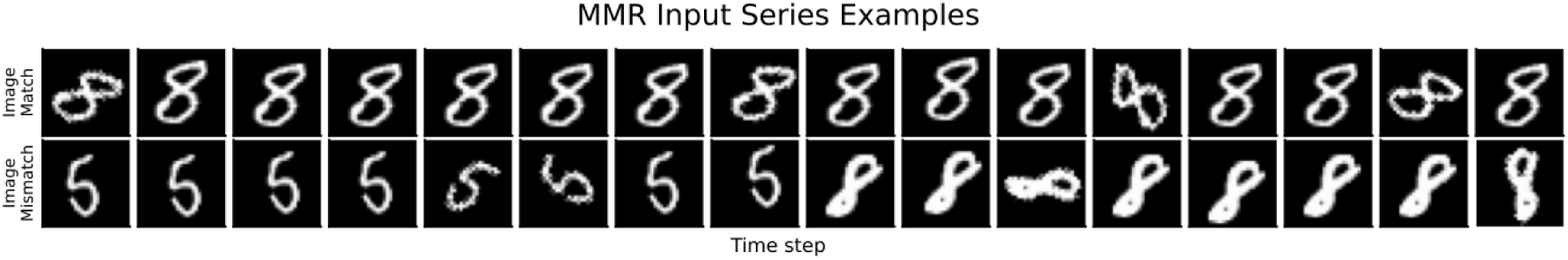
Example image sequences for the matching and mismatching conditions. The top row shows a sequence of MNIST digit images with random shifts and noise, but no changes in the semantic content (matching condition). The bottom row shows a sequence with the same types of visual transformations, but with random switches in the digit identity (mismatching condition).

For networks without activity regularization (see Figure 3; for details see <u>Appendix C1.</u>), most of the PC-inspired conditions showed significant MMR. Specifically, the Contrastive global (T(11998)=−41.58071, p(corrected)<1e-5), Contrastive local (T(11998)=−30.00066, p(corrected)<1e-5), and Predictive local (T(11998)=−40.36738, p(corrected)<1e-5) conditions exhibited significant changes in mean squared layer activity in response to a semantic stimulus switch. The Predictive global condition did not show a significant overall MMR (T(11998)=0.22222, p(corrected)=1). Visual inspection (see Figure 11 in <u>Appendix B2.</u>) revealed that this was due to a positive MMR in the first layer and a negative MMR in the second layer, which canceled each other out in the overall sum (for further discussion, see Discussion 3.1.). Neither the Supervised (T(11998)=2.28285, p(corrected)=1) nor the Untrained (T(11998)=0.72614, p(corrected)=1) conditions showed a significant MMR immediately after the stimulus switch.

**Figure 3:**
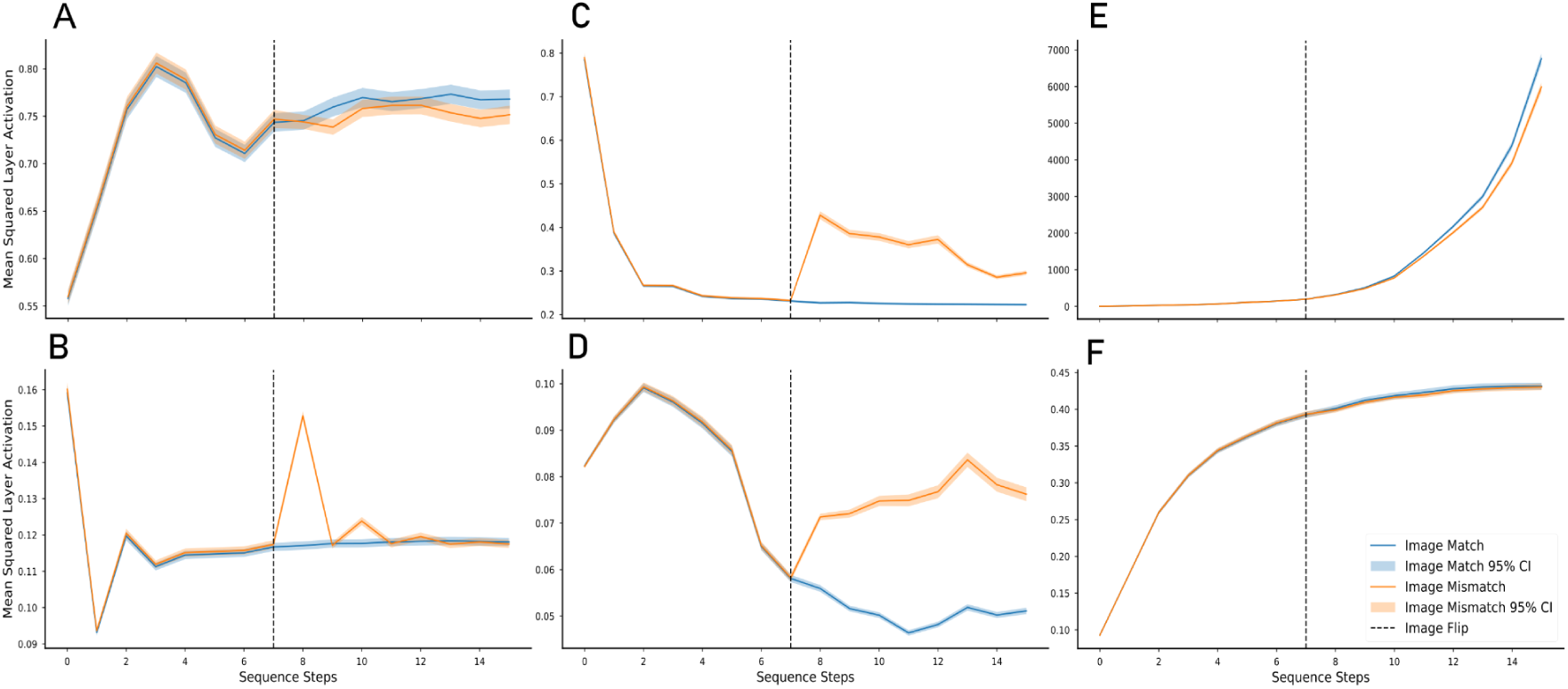
Mismatch responses for unregularized conditions: A. Predictive global; B. Predictive local; C: Contrastive global; D: Contrastive local, E: Supervised; F: Untrained. Each subfigure shows the Mean Squared Average activation of a layer over different time steps, separated for sequences when an unexpected change is happening versus when an expected change is happening. Shaded areas show 95% Confidence intervals.

To investigate whether these effects persist if an additional energy constraint is added, we turned to activity-regularized networks. For activity-regularized networks (see Figure 4; for details see <u>Appendix C1.</u>), all trained conditions, including Predictive global (T(11998)=−11.68170, p(corrected)<1e-5), Predictive local (T(11998)=−20.63438, p(corrected)<1e-5), Contrastive global (T(11998)=−23.97432, p(corrected)<1e-5), and Contrastive local (T(11998)=−15.89864, p(corrected)<1e-5). Even Supervised (T(11998)=17.60957, p(corrected)<1e-5), exhibited a significant MMR, however it is important to note that in this condition the effect was inverted, meaning that the mismatch condition produced less overall activity than the matching condition. The overall effects suggest that activity regularization reinforces or even produces MMR-like effects in artificial neural networks (this effect is further elaborated in Discussion 3.1.). The Untrained condition did not show significant MMR effects (T(11998)=0.72614, p(corrected)<1e-5).

**Figure 4:**
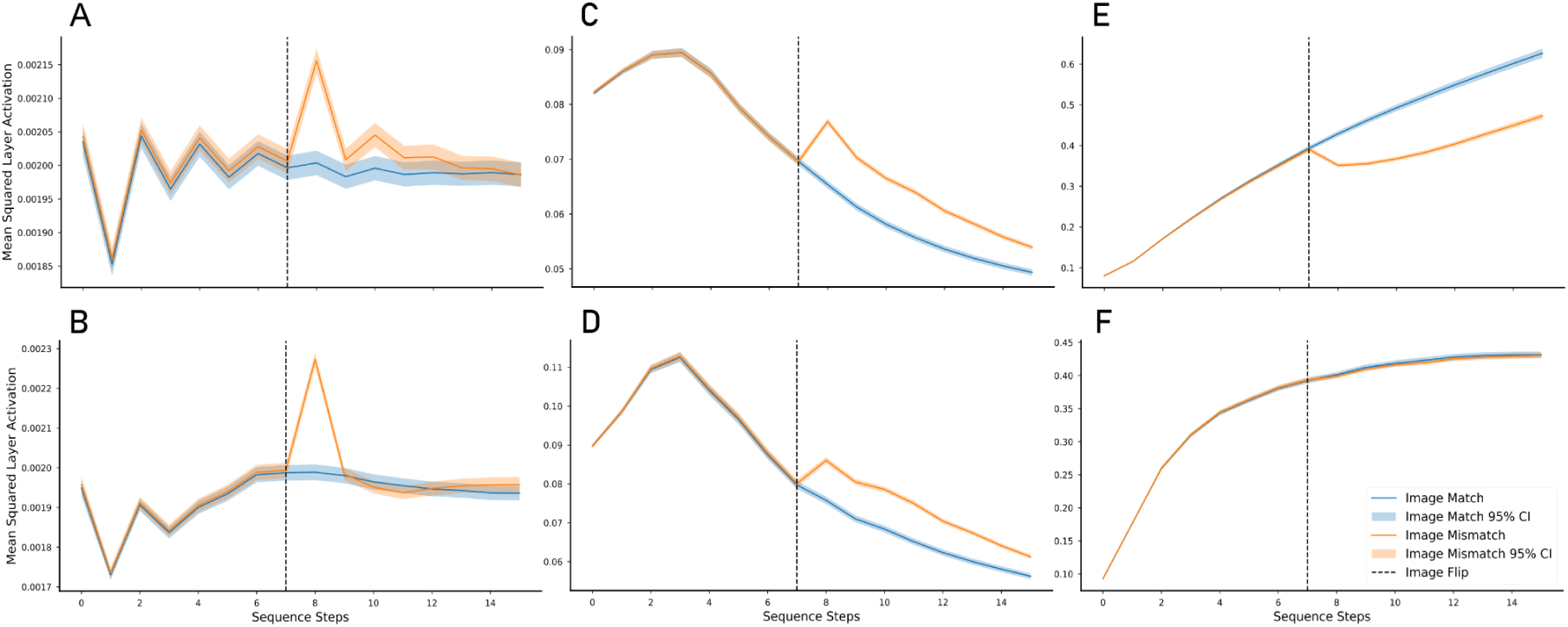
Mismatch responses for activity regularization condition: A. Predictive global; B. Predictive local; C: Contrastive global; D: Contrastive local, E: Supervised; F: Untrained. Each subfigure shows the Mean Squared Average activation of a layer over different time steps, separated for sequences when an unexpected change is happening versus when an expected change is happening. Shaded areas show 95% Confidence intervals.

In summary, the PC-inspired conditions, especially the locally trained ones, were generally able to generate clear mismatch responses, with the exception of the Predictive global condition. Activity regularization appeared to induce MMR-like patterns across all trained models, and even result in (inverted) MMR-like behavior in the Supervised condition.

### 2.2. Prior expectations

Prior predictions are a necessary component of a PC system. They allow the network to match up the prior prediction with incoming information and save energy by only propagating relevant information [54–60]. To measure to what degree the neural network models formed an inherent prior representation, we evaluated the similarities between the neural network’s prior state (negative latent state times recurrent kernel) and the expected future input. If a neural network works according to PC principles, these prior states should start to approximate the future input states over the course of learning. To investigate this effect, we introduced a series of stimuli to all networks and correlated its prior state (latent state multiplied by recurrent kernel) with the actual incoming stimulus. A high correlation in this context means that the prior state approximates the future input well, while a zero correlation means that the prior state and the future incoming input are unrelated (for details, see Methods 4.4.2.).

For unregularized networks, Predictive global (r=0.80507) and Predictive local (r=0.39563) showed clear correlations between the prior and the next stimulus. In contrast, Contrastive global (r=0.01178), Contrastive local (r=−0.01456), Supervised (r=−0.00260), and Untrained (r=−0.00249) did not exhibit any prior-stimulus correlations. Accordingly, only Predictive global (Z(11998)=61.07541, p(corrected)<1e-5) and Predictive local (Z(11998)=23.05043, p(corrected)<1e-5) correlated significantly stronger than the Untrained condition. Supervised (Z(11998)= −0.00612, p(corrected)=1.), Contrastive global (Z(11998)=0.78126, p(corrected)=1.) and Contrastive local (Z(11998)=−0.66089, p(corrected)=1.) did not show prior-stimulus correlations significantly stronger than the untrained condition (see Figure 5; for details see <u>Appendix C2.</u>).

**Figure 5:**
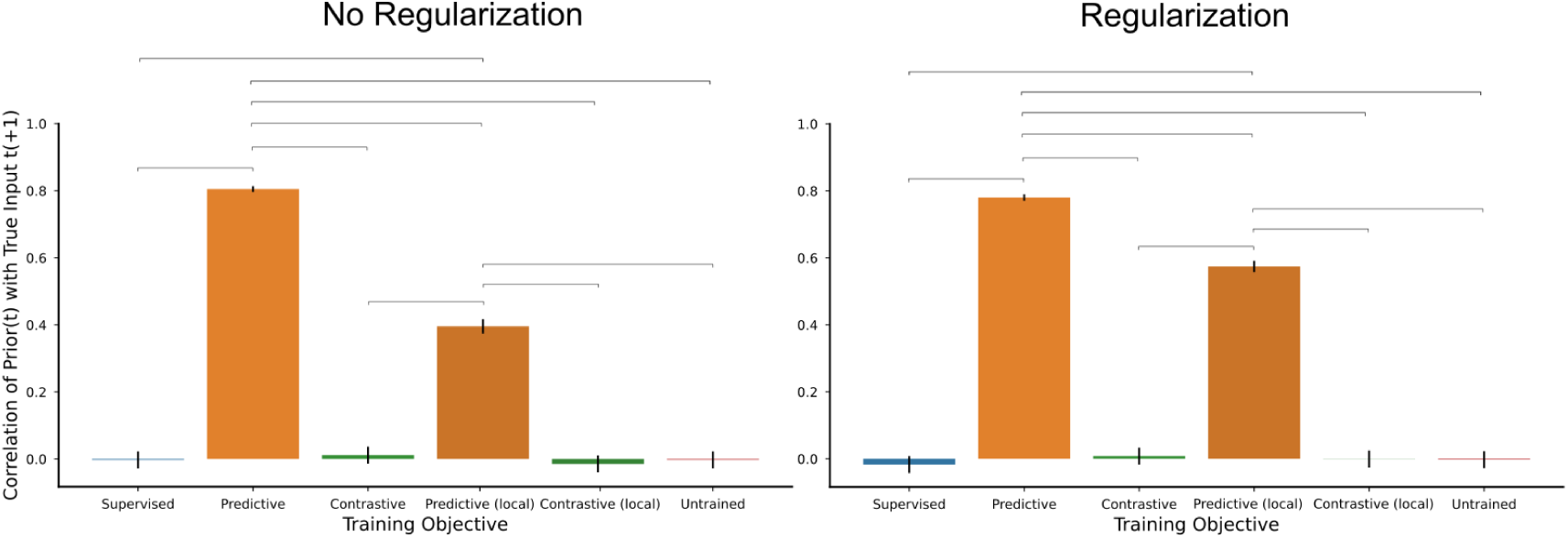
Correlations between priors (project to input level) and future input. Left: No regularization, Right: With activity regularization. High correlation means that the prior (negative recurrent state) projected back to the input level correlates highly with the next (unseen) stimulus in the sequence. Low correlation means the state and the next stimulus are uncorrelated. Solid line: p < 0.0001

The same pattern held true for the activity-regularized networks: Predictive global (r=0.78028) and Predictive local (r=0.57463) demonstrated strong prior-stimulus correlations. Meanwhile, Contrastive global (r=0.00826), Contrastive local (r=−0.00059), Supervised (r=−0.01706), and Untrained (r=−0.00249) did not show any clear prior formation. Like in the unregularized condition, only Predictive global (Z(11998)=57.41855, p(corrected)<1e-5) and Predictive local (Z(11998)=35.97105, p(corrected)<1e-5) correlated significantly stronger than the Untrained condition. Supervised (Z(11998)= −0.79785, p(corrected)=1.), Contrastive global (Z(11998)=0.58876, p(corrected)=1.) and Contrastive local (Z(11998)=0.10407, p(corrected)=1.) did not show prior-stimulus correlations significantly stronger than the untrained condition (see Figure 5; for details see <u>Appendix C2.</u>).

In summary, the Predictive condition was the only one that consistently exhibited the formation of meaningful prior expectations across the different network configurations. The other conditions, including Contrastive, Supervised, and Untrained, did not show evidence of prior formation.

### 2.3. Learned representations

To assess each model’s ability to learn abstract representations, we examined the encoded information content in each model by decoding the original number classes represented by the input from the model’s output state. While the Supervised condition has been explicitly trained to fulfil this task, the PC conditions were optimized on unrelated unsupervised targets. Accordingly, we use better-than-untrained decoding performance as an indicator of the PC network’s capability of learning semantic information under unsupervised learning conditions.

The Predictive global condition (accuracy=0.38667, SE=0.01232) not only performed notably better than the untrained network (T(11998)=7.57907, p(corrected)<1e-5), but not significantly better (T(11998)=2.26123, p(corrected)=0.35644) than the Supervised condition (accuracy=0.36667, SE=0.01219), which still performed notably better than Untrained (T(11998)=5.31126, p(corrected)<1e-5). The Contrastive global condition (accuracy=0.36550, SE=0.01219) also showed substantial learning beyond the Untrained condition (T(11998)=5.17835, p(corrected)<1e-5), though insignificantly lower than Predictive global (T(11998)=2.39393, p(corrected)=0.25026). The Contrastive local condition (accuracy=0.35217, SE=0.01209) (T(11998)=3.65364, p(corrected)=0.00389) as well as the Predictive local condition (accuracy=0.34700, SE=0.01204) (T(11998)= 3.05970, p(corrected)=0.03331) still exhibited significant learning above the Untrained baseline (accuracy=0.32000, SE=0.01180). These results are illustrated in Figure 6 (For details see <u>Appendix C3.</u>).

**Figure 6:**
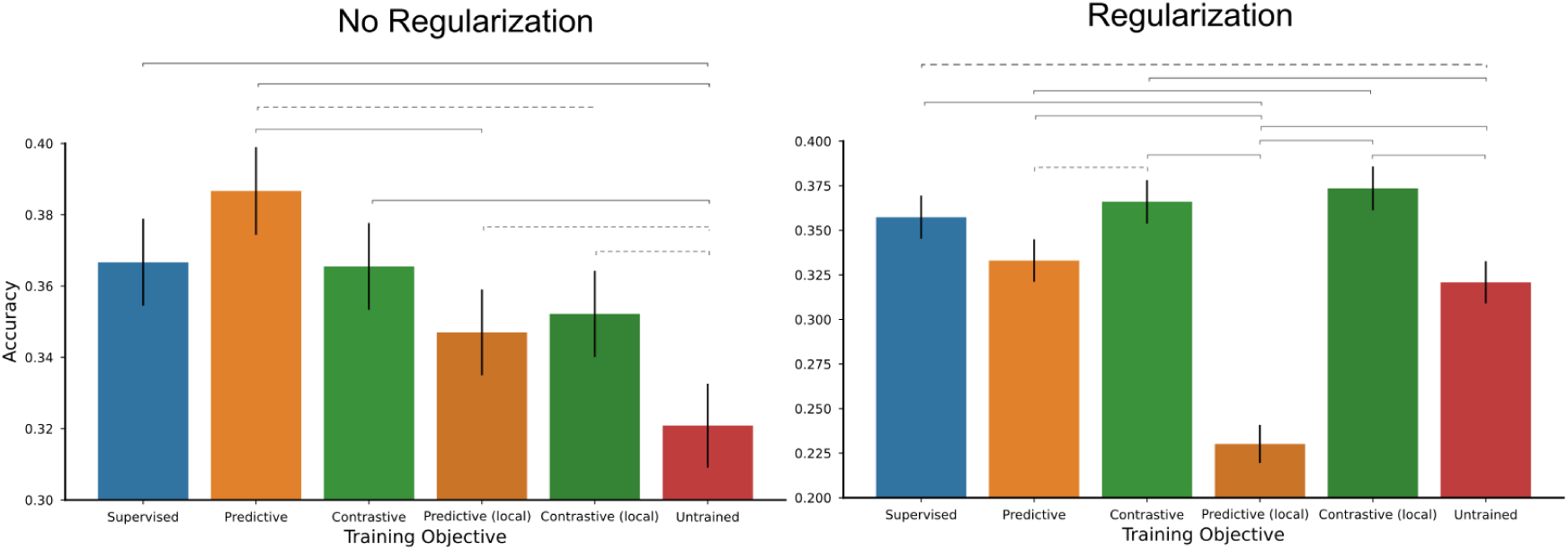
Learning Performance evaluated by classification accuracy of the original stimulus classes based on the output encoding of the neural network models. Left: No regularization, Right: With activity regularization. Dotted line: p < 0.05 Solid line: p < 0.0001

Combining activity regularization with the predictive algorithm resulted in a notable decline of learning performance (see Discussion 3.1.). In this condition, the learning effects diverged more across between models: Predictive global (accuracy=0.33300, SE=0.01192) and Predictive local (accuracy=0.23017, SE=0.01065) both showed decreased learning performance, with Predictive global performing worse than Supervised (T(11998)=2.82368, p(corrected)=0.07133) and on par with an Untrained Network (T(11998)=1.42093, p(corrected)=1.) Predictive local even performing notably worse than the Untrained condition (T(11998)=−12.62837, p(corrected)<1e-5). In contrast, Contrastive global (accuracy=0.36600, SE=0.01219)(T(11998)=5.23532, p(corrected)<1e-5) and Contrastive local (accuracy=0.37350, SE=0.01224)(T(11998)=6.08825, p(corrected)<1e-5) exhibited increased learning compared with the Untrained condition. The Supervised condition still performed notably better than the untrained condition (T(11998)=4.24578, p(corrected)=0.00033). These results are illustrated in Figure 6 (For details see <u>Appendix C3.</u>).

To conclude, our analysis of semantic learning effects showed that all algorithms in almost all settings exhibited significant learning of category information beyond the untrained baseline condition. In the unregularized setting, all training conditions showed significant learning effects beyond the untrained baseline. While this is to be expected for the Supervised condition, it shows that the PC objectives as well are capable of instilling semantic information into a model despite their unsupervised training objective.

### 2.4. Gain control

In PC, the noise level of the output signal is correlated with the noise level of the input. Gain control is the ability to manipulate the output noise level independently of the input noise [40,41]. As gain control is an important mechanism in PC literature [40], we investigated whether it is possible to simulate this mechanism in PC networks using weight regularization. We created two input conditions with high and low noise levels and evaluated whether weight regularized networks significantly reduce the difference in noise level of the model’s output. We did this by comparing the variance ratio between low and high noise input for unregularized versus weight regularized networks.

Our results showed that weight regularization results in a gain control-like effect, reducing the neural activity’s variance ratio between high noise and low noise input stimuli. Contrastive (Z=−67.08483, p(corrected)<1e-5), Predictive (Z=−64.67718, p(corrected)<1e-5), and Supervised (Z=−67.08477, p(corrected)<1e-5) all showed significant reduction in output variance ratios between high noise and low noise when weight regularized. From visual inspection of Figure 7, it can be seen that all algorithms have undergone notable reduction in variance differences with applied weight regularization. However, weight regularization did not lead to perfectly identical output variance distributions for any algorithm, with Contrastive (Z=−67.08478, p(corrected)<1e-5), Predictive (Z=−66.92948, p(corrected)<1e-5), and Supervised (Z=−36.19967, p(corrected)<1e-5) all still showing significance variance difference between high noise and low noise samples (for details see <u>Appendix C4.</u>).

**Figure 7:**
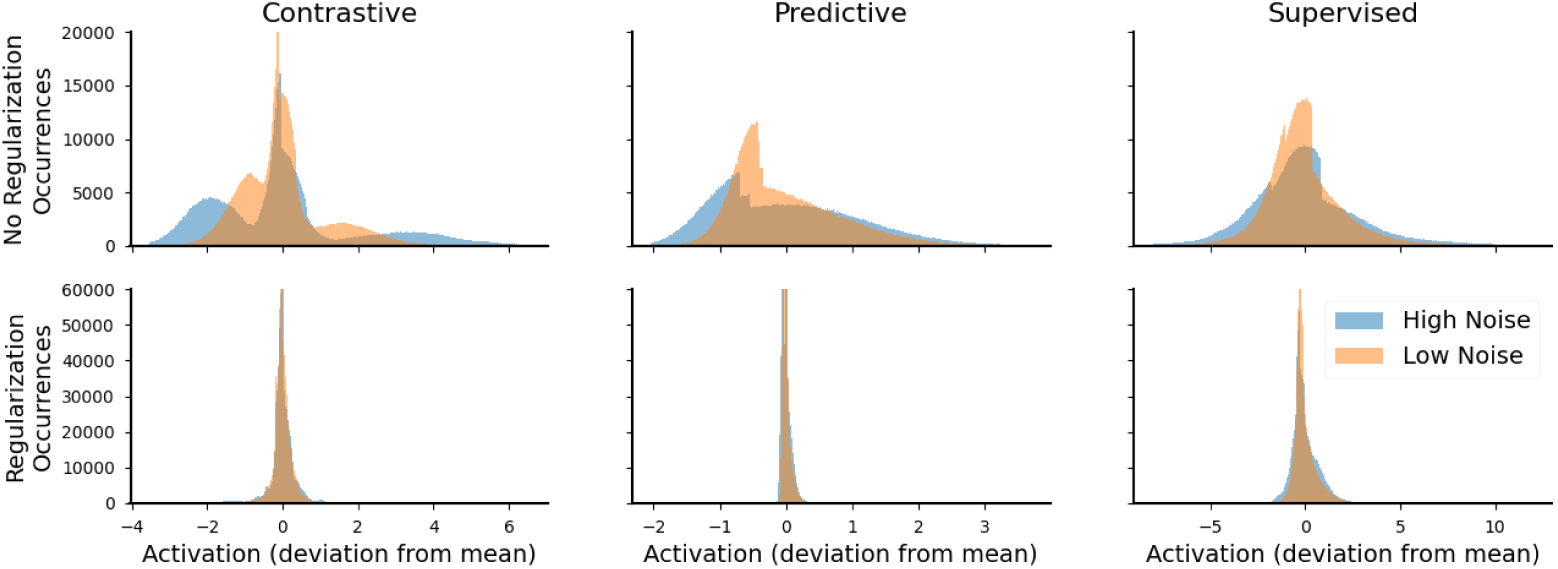
Distributions of output activations for unregularized and weight-regularized networks, for high noise and low noise inputs. Output activations from high noise inputs are shown in blue, output activations from low noise inputs are shown in orange. Weight regularization leads not only to normalization of the neural output distribution, but also reduces the ratio in output variance in response to high noise and low noise inputs.

## 3. Discussion

The overarching aim of this research was to investigate whether common neural network architectures can be adapted to exhibit key signatures of PC. Specifically, we were interested in exploring the models’ ability to form prior expectations, generate mismatch responses, and learn robust representations in an unsupervised manner. The results of this study suggest several key insights: (1) Networks trained with predictive coding-inspired objectives generally exhibited more characteristics of predictive coding compared to non-PC-inspired objectives. (2) The local Predictive condition (without activity regularization) was the most likely candidate for predictive-coding like effects due to showing clear MMR, priors, and learning effects. (3) The contrastive approach showed very pronounced mismatch responses and good learning performance, especially with activity regularization. However, it did not lead to the explicit formation of prior expectations. (4) Activity regularization seemed to evoke mismatch-like effects across all trained networks, but decreased the learning performance of the predictive and locally predictive networks. (5) Weight regularization assimilates output variance of networks, irrespective of the noise level of the input. This mechanism can be used to model and investigate the biological process of gain control.

In the following section, we discuss these findings in more detail.

### 3.1. To what extent did the PC inspired models exhibit landmarks of biological PC?

#### Mismatch Responses

All PC-inspired networks showed some form of MMR (i.e. stimulus mismatch induced positive activity) in at least one layer. We found the strongest MMR in the Contrastive condition, likely due to the fact that the Forward-Forward implemented here is directly optimized towards maximizing the output activity after occurrence of mismatching stimuli. The Predictive local condition also evoked clear MMR. Interestingly, the Predictive global condition produced a negative MMR in the first layer and a positive MMR in the second layer, which cancelled out when added together (see Figure 11 in <u>Appendix B2.</u>). This phenomenon likely arises from the fact that the globally defined predictive function is defined to facilitate MMR at the final layer, which might lead to opposite effects in the previous layers, meaning that (goal-oriented but negative) weight patterns that are punished in the final layer are more likely to appear in previous layers. A comparable pattern is observed in biological neural networks as well, where repetition suppression and repetition enhancement can occur at different levels of the cortex [41]. However, further research is needed to determine if it is analogous to neurophysiological MMR or an unrelated occurrence in the artificial network. Overall, the clear MMR spikes in PC-inspired networks sets them apart from Supervised networks, which does not show such clear spikes. These results provide further evidence that MMR are a consequence of prediction-based learning, as claimed in PC literature [51]. They further confirm that complex semantic information can be learned in an unsupervised fashion by maximizing the contrast in MMR (as proposed in [45]).

MMR responses were strongly enhanced or even evoked by activity regularization. This suggests activity regularization may be a promising approximation of the energy-saving principle behind PC [50]. Interestingly, we find that activity regularization evokes an inverted MMR-like effect in Supervised recurrent networks, where the output activity for mismatching stimuli is lower than for matching stimuli. This does not directly align with the idea of PC, where energy should be saved by only propagating unexpected information, meaning that mismatch activity is expected to be larger than expected activity. This effect can be explained by the combination of the Supervised learning objective in combination with activity regularization: Supervised RNN models have the tendency to build up activity over time through accumulating activation over the recurrent state (this is an issue related to the exploding gradients problem and discussed in [81,82]). In unregularized networks, this leads to an exponential increase in activity (see Figure 3) as long as the network is presented with unchanching patterns that accumulate and amplify in the recurrent state. Now when a mismatching input appears, the activity builds up slightly slower because the new input tends to cancel out activity rather than further amplifying existing patterns. In the activity regularization setting, this activity accumulation is decreased due to increased penalization of high activity (see Figure 4). However, similar to the unregularized condition, the stimulus switch tends to cancel out activity rather than amplifying existing patterns, leading to reduced activity for the mismatch condition. This might explain the effect of the MMR-like pattern in the Supervised model with activation regularization.

Further, we found that the application of activity regularization in combination with a Supervised objective, resulted in unexpected behavior. The Predictive condition performed well overall, but was heavily impaired when combined with activity regularization. An interesting aspect of this phenomenon is that the Predictive models with activity regularization still exhibited very strong correlations between the current prior and the future state, but seemed to encode drastically less class-specific semantic information. This suggests that the activity regularization forced the loss in the Predictive objective to only focus on the Predictive information which is shared between classes, creating relatively high predictions from this generally shared variance, while at the same time cutting all the activity that would carry important information for specific classes, but which occur less often and therefore contribute too little signal to withstand the activity regularization.

#### Prior expectations

Contrary to the broad occurrence of MMR, only the Predictive condition showed clear prior formation, while the Contrastive, Supervised, and Untrained conditions did not. Empirical evidence shows that predictions in the brain are usually organized topographically. This has been widely shown in the visual, auditory, sensory, and other systems [62–67]. As PC builds upon this idea of neuron-by-neuron topographically ordered predictions, this suggests that a population-activity based contrastive model such as Forward-Forward [45] might not be a perfectly suitable model of PC. Further, this suggests that explicit definition of a prior in the Predictive algorithm may lead to the appearance of an MMR as a secondary effect, while the opposite (i.e. the explicit definition of a MMR in the Contrastive algorithm implying the formation of a topographically organized prediction) does not hold. This insight can be further used to investigate alternative models of PC and Bayesian brain theories [42,43,51,68–71]. MMR are among the main phenomena that lead to the development of the theory of PC and are frequently used as evidence for it [41,51]. If there are learning systems that exhibit MMR, but do not exhibit explicit prior predictions, then models such as the ones presented here can help in investigating the theory by evaluating whether a predictive MMR model or a non-predictive MMR model fits the brain better.

#### Learned Representations

Finally, to assess the learning effects of the different training approaches, we compared the performance of the models on the task of classifying the original MNIST digit classes based on the encoded representations in the networks. In general, predictive and contrastive models showed better-than-random learning effects. Supervised showed the strongest learning effects, because the algorithm was specifically optimized to perform a classification task on a similar set of input stimuli. One important aspect to consider is that the Predictive model used in this study performed well at encoding semantic information about stimuli, while similar investigation into the classical PredNet architecture [72] did not show strong encoding capabilities. This is possibly related to the lack of semantic encoding occurring in the convolution-based PredNet architecture (see Figure 8), which has been fixed in the presented predictive model. While the Supervised algorithm still showed the best learning performance, our results show that PC-inspired algorithms might be useful for many learning contexts, adding to other research confirming PC’s usefulness for tasks like transfer learning and generalization [73].

**Figure 8:**
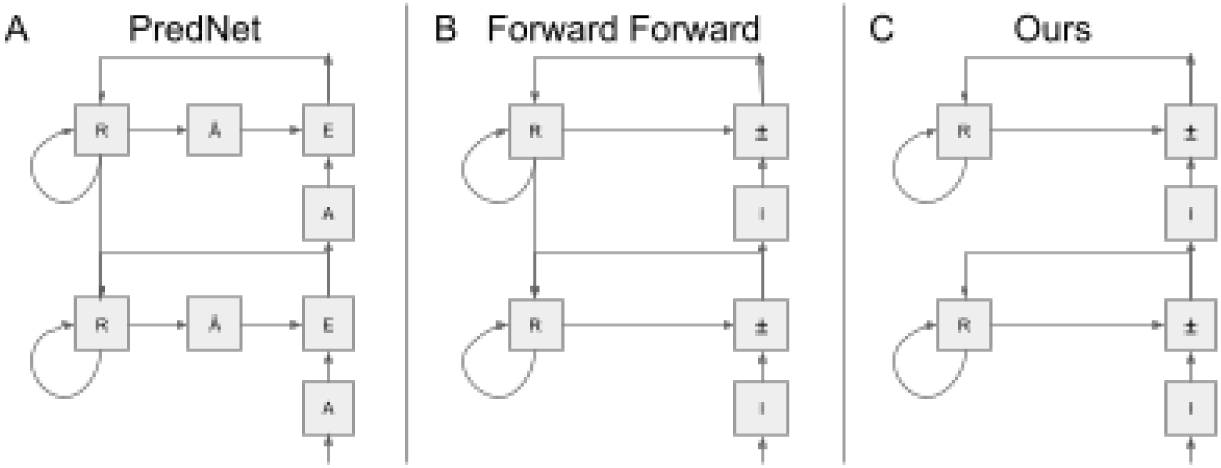
A. Illustration of the PredNet Architecture [49]. B. Illustration of the Recurrent version of the Forward Forward architecture [45]. C. Illustration of the Simple RNN Architecture used here. While PredNet is a high level approach for self-supervised video learning, we use SimpleRNN to create the most minimalistic and assumption-free type of network capable of expressing prior predictions. There are 3 main differences between the two models: (1) The top-down connections presented in PredNet are omitted in the SimpleRNN for the sake of simplicity. (2) The layers in PredNet are specified for learning of visual information, using convolutional layers, convolutional LSTMs, max-pooling layers. In the Simple RNN, we use only a minimal amount of weights, consisting of one input kernel and one recurrent kernel. (3) PredNet maintains the pixel-level state of the image throughout the entire network hierarchy, meaning that the network does not encode the image sequence, but only applies changes to it. This means that PredNet output states do not provide a biologically plausible representation of neurons that might be interpreted as pure positive firing patterns of neurons. This maintenance of the pixel-level variable in combination with using ReLU activation functions (which are only capable of expressing positive values) also requires an explicit handling and splitting up of positive and negative correction errors. In the SimpleRNN model, the states are encoded and maintained as purely positive values (interpretable as firing patterns). In this network, the expression of positive and negative error corrections takes place through positive and negative weights in the input and recurrent kernel, which are applied *before* the activation function is applied. The principle of Forward Forward on the other hand is more flexible than PredNet. In the original paper [45], the algorithm is more defined as a target objective rather than an architecture, and can be implemented in a multitude of different architectures, recurrent or nonrecurrent. The architecture detailed here is an approximation of the recurrent example of the Forward Forward algorithm presented in the original paper.

#### Gain Control

In an additional analysis we found that weight regularization added to neural networks during the training process results in a variance normalization of the output activations. In unregularized networks, the variance of the network’s output activity correlates with the variance of the input. However, when weight regularization is applied, this correlation is weakened, meaning that noisy inputs almost produce the same output activity variance as clean inputs. We successfully showed that weight regularization can be used to implement gain control mechanisms in DNN models of the brain. Such a gain control mechanism may achieve two aims, consistent with the PC framework: 1. scaling output variance to compensate for input variance, and 2. encouraging the network to learn sparse representations and minimize activity to conserve energy [37,39,74]. Additionally, this effect should be taken into account when using DNN as neuroconnectionist models while changing the noise levels in the input (such as [75,76]). In such cases, networks trained using regularization could lead to different distributional properties of the activation, which might affect outcome or decodability in downstream (encoding, decoding, RSA) analyses.

#### Summary

Taken together, these results indicate that under certain circumstances, the neural network models trained with PC-inspired objectives were able to display hallmarks of predictive processing. Specifically, the locally trained predictive objective (without activity regularization) in a RNN exhibited these traits most closely. This model was able to fulfill the expected landmarks of PC, including the formation of meaningful priors, the generation of mismatch responses, and the learning of informative representations, without requiring extensive gradient propagation throughout the network hierarchy. This strongly supports the idea that simple predictive objectives can serve as valid approximations of PC. These models can exhibit behaviors resembling information processing in biological neural networks better than standard DNN models. Our results not only show that PC is effective as a learning mechanism (which has been demonstrated in other studies; [11,44–47,77–80]), but also connect the learning effects to the phenomenological markers theorized as part of PC theory (such as priors and MMR), confirming them as a possible and logically coherent consequence of the PC mechanism.

### 3.2. What are the key limitations of the current study and how can they be addressed?

While this study provides valuable insights into the relationship between artificial and biological learning, it highlights the need for further exploration and refinement of PC-inspired neural network models. Future research should scale up the model complexity by exploring more sophisticated architectures with additional layers and increased capacity. This could uncover new emergent properties and better align the artificial networks with the brain’s hierarchical structure. Specifically, incorporating deeper networks with longer propagation of backwards errors could better mimic the hierarchical information processing theorized by PC.

Another possible improvement is to test these algorithms on more challenging and diverse tasks beyond the simplified moving MNIST paradigm used in this work. The PredNet model [49], for example, has been evaluated on high level visual data including more complex and naturalistic inputs [83]. However, PredNet has been criticized for issues related to its biological plausibility [72]. The predictive model used in the current study aims to address these biological plausibility concerns, but it has not yet been tested on larger-scale tasks. Similarly, the contrastive model, which is based on the Forward-Forward algorithm [45], as well as other related algorithms [79], have only been evaluated on MNIST or moving MNIST, which are relatively simplified tasks. MNIST or moving MNIST may not generalize well to more complex computer vision tasks [84]. By testing these PC-inspired models on a wider range of tasks, including those with more naturalistic and diverse inputs, researchers can better assess their generalization capabilities and further explore the computational principles underlying biological neural information processing.

### 3.3. Implications for biological and artificial neural learning

The insights from this research can be used to create better DNN models of the brain. The neuroconnectionist framework relies on creating neural network models and experimentally altering different mechanisms to investigate how changes in the model improve or decrease its similarity to the brain [1,16]. However, research into PC often relies on biologically implausible machine learning networks [20,72,85] despite previous research showing that well-performing machine learning models are not always better neuroconnectionist brain models [1,24]. The results presented here suggest that the PC-inspired models, while currently only sparsely being used as neuroconnectionist research models [44], might be a promising future direction in brain modelling.

From a machine learning perspective, the results from this paper support existing evidence that PC-inspired algorithms based on Predictive as well as Contrastive objectives [45,49] are promising methods for building effective learning algorithms. The models presented in this work perform well in their role as unsupervised machine learning models, even after addressing many of the biologically implausible aspects that have been criticized in classical deep neural networks. Specifically, such models could serve as a foundation for developing new artificial neural network models for time-dissolved or sequential tasks such as time series or video prediction. While PC-inspired models currently may not surpass existing state-of-the-art models in unsupervised learning [86–88], the elegant way in which PC integrates a prediction-based framework with complex semantic learning systems poses a strong foundation for creating simple and effective theory-driven models for unsupervised learning.

## 4. Methods

### 4.1. General Model architecture

To implement the PC-inspired neural network models, and to facilitate comparisons between distinct models, we used a simple RNN architecture as a basis for all models. We chose this straightforward RNN architecture to demonstrate the core properties of PC in a maximally simple and well-controlled setting. Specifically, all conditions were trained on a model with two vanilla RNN layers - both composed of a forward kernel as well as a recurrent kernel. The recurrent connections in the RNN allow the model to generate some form of prior expectations, which can then be integrated with incoming sensory input. Specific details on the architecture and hyperparameters are given in <u>Appendix A</u>. All code to replicate this study will be published under github.com/DiGyt/predictive_coding_algorithms.

By using a simple RNN rather than more complex architectures, we aimed to isolate the key PC principles without the confounding factors that could arise in deeper or more specialized network designs [89]. Further, this constant architecture allows us to isolate training effects induced by the training objective and optimization procedure, omitting effects created by varying architectures.

### 4.2. Training environment

The neural network models in this study were trained on a moving MNIST [90] paradigm, which was specifically designed to present the models with a challenging yet controlled set of input stimuli. This paradigm involves a series of MNIST digit images that move across the input field in a particular order, with the digits moving at different speeds (comparable to [79]) (see Figure 9). Specifically, the movement of the MNIST digits follows sine wave patterns, with the phase and direction of the sine waves on the horizontal and vertical axes determined by the class of the number being presented. This approach encourages the networks to develop robust representations that link the digit semantics to their dynamic visual features, rather than just memorizing static pixel-level details. We use MNIST because it is a typical dataset to test new DNN architectures on, and has been used as a test dataset for various novel DNN architectures [45,79]. However, as a dataset for semantic classification, MNIST is notoriously easy to solve [91]. In order to enhance classification difficulty and improve generalization, we applied a variety of random augmentations to the input images. These augmentations included shifts (up to ~±21% of the receptive field with a standard deviation of ~14% of the receptive field) and the addition of Gaussian noise (+33% standard deviation relative to noise-free images). By introducing these transformations, we ensured that the networks could not simply memorize the pixel-level details of the inputs, but had to learn more abstract representations. Additionally, all input images were normalized to a 0 to 1 scale, rather than the original 0 to 255 range, to help standardize the data.

**Figure 9:**
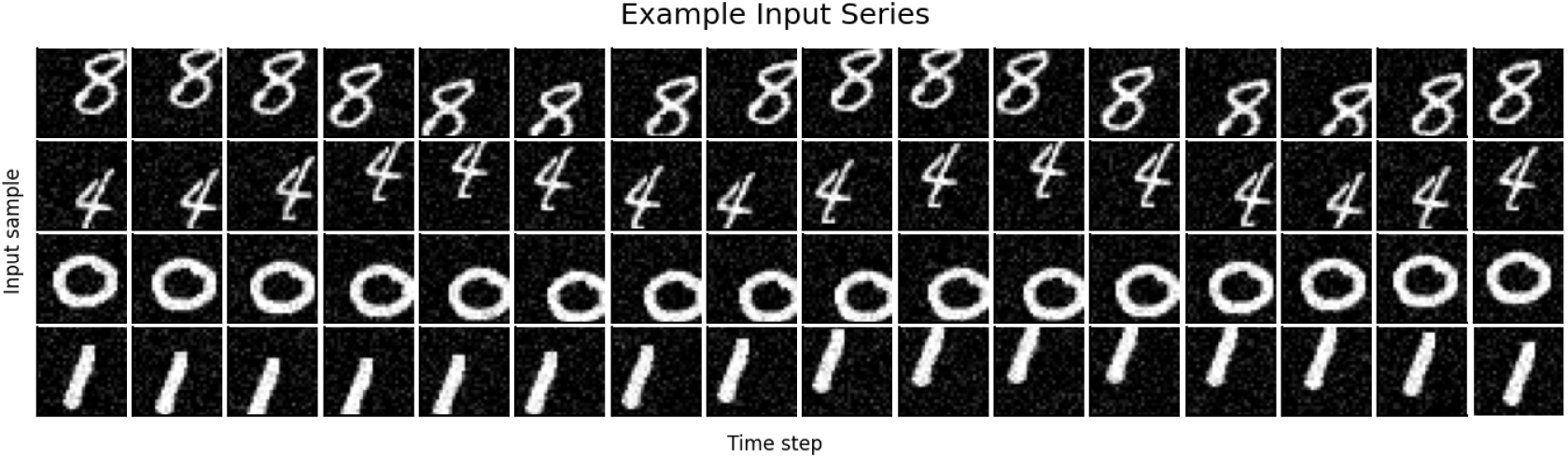
Illustration of the input series. Networks were trained according to a moving MNIST paradigm. For each batch, multiple samples were fed to the network. Each sample contained a series of images, depicting a number moving across the visual field. Each separate number class moves across the visual field according to specific rules (direction and magnitude in change of x and y coordinates per time step). These rules were shared between samples of the same number class, but differed from samples of other number classes.

The training dataset consisted of 54,000 MNIST samples drawn from the original training set, with an additional 6,000 images held out for testing. During training, the data was shuffled each epoch and then batched up in batches of 512 samples per batch (and 240 samples for the final batch).

### 4.3. Model variations

#### 4.3.1. Training objectives

In order to easily compare different predictive-coding like training objectives with supervised backpropagation while omitting potential nuisance factors from the network architectures, we adapted two common PC-inspired algorithms to work with a simple RNN architecture. This is especially relevant since common PC algorithms such as PredNet are specifically designed to work with Image-type data (including convolutional network layers) and therefore comparisons to other algorithms on a fundamental level are not trivial. In this section, we describe the core inspiration for these algorithms and how we adapted them. The different training conditions are illustrated in Figure 1.

##### Predictive

One of the training procedures we investigated was a predictive, or autoregressive, objective. This approach was inspired by the PredNet model, which aims to predict the next frame in a sequence of inputs. PredNet is a predictive-coding inspired DNN architecture published by Lotter et al. [49]. The architecture consists of a series of stacked modules, each comprising a convolutional input layer, a recurrent layer, a prediction layer, and an error representation layer. Each module makes local predictions about the input, and the deviations from these predictions are forwarded to higher layers, allowing the network to learn the temporal structure of visual data. This hierarchical approach enables the model to predict future frames in video sequences while capturing essential aspects of object movement and scene dynamics, meaning the loss is simply the mean squared error (MSE) between the model’s final output and the stimulus image of the next time step. However with its architecture, PredNet is explicitly designed to create pixel level predictions as their output. Due to the fact that the objective of PredNet is to approximate the output representations as close as possible to the future input, the output representations of PredNet are necessarily forced to retain the same level of representation as the input, without encoding the input to more condensed representations, as it would usually happen in image recognition DNN (including typical CNN architectures). This means that PredNet is limited to operating on image-like data and performing semantic predictions at the sensory input level, rather than learning more abstract, encoded representations.

To solve this problem and to make the predictive objective more generally applicable to a wider range of neural network architectures, we implemented an alternative version of the predictive training approach. In our implementation, the loss function consisted of a mean squared error (MSE) loss between the current “prior” (the previous hidden state multiplied by the recurrent kernel and projected back through the inverse input kernel) and the actual input stimulus one step into the future. This change allows us to encode representations into an arbitrary set of neurons, rather than maintaining the representations in the input space like PredNet does. Further, this loss is adaptable to various DNN model architectures, allowing us to implement the predictive objective in the more comparable setting of a simple RNN architecture.

By using this predictive, autoregressive objective, we encourage the neural network models to learn to generate predictions about upcoming inputs and update their internal representations in a way that they approximate the future input as close as possible given the current information. This training procedure aligns with the key principles of PC, where the brain is thought to constantly generate predictions about sensory inputs and update its models based on the mismatch between predictions and observations. Implementing this predictive objective in a more general way, rather than relying on specialized architectures, allowed us to explore the potential of this approach across a broader range of neural network models.

##### Contrastive

Another training procedure we investigated was a contrastive objective, inspired by the Forward-Forward algorithm proposed by Hinton [45]. The Forward-Forward algorithm works as a type of contrastive optimization loss, where an arbitrary network learns about stimuli by being able to separate valid (positive) data from invalid (negative) data. This happens on a layer-wise level, such that for example layer-wise activity is minimized for valid data, and maximized for invalid data. In this setting, the algorithm is comparable to the general concept of PC, where expected (valid) stimuli should evoke no activity, while unexpected (invalid) stimuli evoke maximum activity. In the original paper, a broad concept for a multi-hierarchical implementation of the algorithm is presented, however, as a preliminary investigation, the paper presents only working examples of more simple, feedforward context, where the network effectively performs contrastive objective, separating coherent input stimuli from incoherent ones. This basic form of the model was also implemented here, to facilitate comparisons with PredNet.

The network architecture for this condition was derived from Hinton’s original description. The loss function was defined to minimize a layer’s activity when the next input image moves as expected (e.g., the numbers move according to predictable rules), and to maximize activity for stimuli that are unexpected (e.g. the numbers move according to non-predictable rules). In contrast to the predictive objective, this does not happen on a neuron-by-neuron level, where the resulting neuronal activity is expected to match activational patterns that predict the upcoming stimulus, but instead only considers the overall activity of the layer, which should be minimized for expected stimuli and maximized for unexpected stimuli. To implement this contrastive objective, the input data was split into positive samples (expected stimuli) and negative samples (unexpected stimuli). The positive stimuli consisted of normal number sequences that would make the MNIST numbers move as expected, while the negative stimuli included random semantic number switches, resulting in numbers that are not aligned with the rules according to which they should move. By implementing this contrastive objective, we aimed to encourage the neural network models to learn representations that are sensitive to expected versus unexpected stimuli, a key characteristic of PC. This training procedure provides an alternative approach to the predictive objective, allowing us to explore different ways of instilling PC-like principles into the models.

##### Supervised

As a first control condition, we used a standard supervised backpropagation approach. In this case, the target objective for the neural network models was to classify the image class of the current number shown in the input sequence. The model architecture for this condition is identical to the other models, with the only exception that, for this model, the final layer output is mapped onto a one-hot representation of the stimulus classes, allowing to train the network in a supervised fashion.

The use of supervised backpropagation provides an important baseline for comparison against the PC-inspired training procedures. Supervised learning, which relies on classical deep learning optimization rules like stochastic gradient descent and backpropagation, is known to be effective at training artificial neural networks to perform a wide range of tasks.

However, these supervised training approaches are often defined based on information-theoretical ideas rather than implementing biologically plausible objectives. By including a supervised backpropagation condition, we can assess how the performance and characteristics of the PC-inspired models compare to a standard, widely-used training method that is not explicitly designed to capture the computational principles of biological neural learning.

##### Untrained

As another baseline control condition, we used a model initialized of the same architecture as all the other models, but without undergoing any training process. Comparing the results of the trained models to this untrained random network baseline will help us determine the specific contributions of the different training objectives, ensuring that any observed signatures of PC are indeed a result of the learning process, rather than simply inherent in the recurrent model architecture itself.

#### 4.3.2. Localized optimization procedure

In backpropagation, the error gradients are propagated through a long nested chain of partial derivatives that are all directed towards optimizing the output of the final layer [10]. A common critique on the biological plausibility of backpropagation is the implausibility of conveying such learning gradients backwards through a high number of biological neural layers [3,11,15]. In contrast to that, learning signals in the brain are usually transmitted using chemical or electrical signals [92–95], which are too diffuse and imprecise to transmit exact gradients over long hierarchies of layers. While it is theoretically possible to send gradient-like learning signals through a long hierarchy of neural layers [15], this becomes more unlikely, the longer the hierarchy becomes [3,11–15]. Accordingly, the biological brain is widely believed to use local learning gradients [7,11,12]. Importantly, both the predictive and the contrastive training conditions can be implemented in a localized, layer-by-layer fashion rather than using an end-to-end backpropagation approach throughout the network. For the contrastive condition, this localized training is possible because of the use of positive versus negative samples. The contrast in firing patterns can be learned at every level of the hierarchy, allowing a training procedure that first creates maximum contrast in output activity in the first layer, then the second layer, and so on. Similarly, the predictive objective can also be applied in a localized manner. By approximating the current “prior” (the previous hidden state projected back to the next lower level) to the next input state, this predictive loss can be computed at any given level of encoding. This allows for a training procedure that enables learning the first layer, then the second layer, and so on, without the need for full end-to-end backpropagation.

Accordingly, we introduced additional localized conditions: besides the Predictive/Contrastive global conditions, which predict/contrast only at the last layer and use backpropagation for propagating the gradients through the network, we additionally introduce a Predictive/Contrastive local condition, where gradients are only propagated within one layer. By exploring these localized training procedures, we aim to investigate whether PC-inspired objectives can be effectively implemented in a more brain-aligned fashion, without relying on the biologically implausible backpropagation algorithm.

#### 4.3.3. Activity regularization

One key hyperparameter we investigated was the use of regularization. Specifically, we compared the results of the models trained with and without activity regularization. Activity regularization enforces sparsity in the layer activations, which provides an interesting analogy to the energy-saving principle behind PC [50,96].The rationale for exploring activity regularization is that it can serve as an approximation to one of the underlying assumptions of PC, namely that the brain is thought to minimize the energy required for information and only propagate the residual, or “prediction error” information [97]. Similarly, activity regularization encourages the neural network models to learn sparse, efficient representations which minimize the activity required to represent stimuli.

By comparing the performance of the models trained with and without activity regularization, we aimed to gain insights into the role of this type of regularization as a proxy for the computational mechanisms underlying PC. This analysis allowed us to explore whether the incorporation of this energy-saving principle, even in a simplified form, can lead to neural network models that better exhibit the hallmarks of predictive processing in the brain.

### 4.4. Model evaluation

We selected three specific evaluation metrics - mismatch responses, prior expectations, and learned representations - to align with the key behaviors we would expect from a neural network model exhibiting the principles of PC. In the following, we describe how each of these metrics are quantified.

#### 4.4.1. Mismatch responses

In PC, the occurrence of MMR is directly tied to deviations from what was expected, resulting in elevated activity [61]. In neuroscientific literature, mismatch responses are a widely observed phenomenon, occurring in various modalities [53,98,99]. We expect similar MMR patterns in PC-inspired networks. To assess the models’ ability to detect and respond to unexpected or deviant stimuli, we evaluated their MMR (measured by deviations in the evoked neural response) to a series of matching and mismatching input sequences (see Figure 2).

The matching series condition consisted of a sequence of MNIST images with random shifts and rotations, but without any semantic changes to the digit content. In contrast, the mismatching series condition included shifts, rotations, and random switches in the semantic content of the images (e.g., a 3 suddenly changing to a 7). We ensured that the augmentation between match and mismatch condition did not show any base level differences that might result in a difference in mean-squared activation from the network’s layers. We performed a Kolmogorov-Smirnov (D=0.000, p=1.000) on the pixel-level distribution of the two image conditions, to make sure that the core statistics of the matching group does not differ significantly from the mismatching group. We then measured the mean square layer activations of the networks over time, approximating evoked responses for the matching and mismatching conditions. Our metric of interest was whether the model output produced significant deviations between the match and the mismatch condition for the input immediately following the stimulus switch. This deviation was compared using independent sample t-tests, corrected for multiple comparisons across models using the Bonferroni correction. By analyzing the mismatch responses of the neural network models, we aimed to assess their ability to detect and respond to prediction errors, a key characteristic of PC. The comparison of the matching and mismatching conditions allowed us to isolate the models’ sensitivity to unexpected changes in the input stimuli. As a metric for inference we evaluate the difference between the matching and the mismatching condition, immediately following a stimulus switch. We performed undirected independent t-tests with significance levels corrected for each time step and each condition using a Bonferroni correction.

#### 4.4.2. Prior expectations

Prior predictions are a necessary precondition for a PC system, allowing to match up the prior prediction with incoming information, to save energy by only propagating relevant information [54–60]. Priors in the DNN context mean that there exists a representation of what input values the network expects to come up in the next step, which is then matched up and integrated with the actual incoming stimulus. If a neural network works according to PC principles, these prior states should start to approximate the input states over the course of learning. This would mean that the neural network forms expectations which not only convey information to the next state, but explicitly model what will happen in the next state (in order to implement expectation-related suppression of predicted stimuli [41,100]).

To measure to what degree the neural network models formed an inherent prior representation, we evaluated the similarities between the neural network’s prior state (negative latent state times recurrent kernel) and the expected future input. We assessed the prior formation by projecting the prior state back through the network to the input level. This projected prior was then correlated with the future input to express the similarity between the prior and the future incoming input. As a second confirmatory analysis, we also performed this analysis at the encoded level by propagating the future inputs through the network in a forward direction and correlating the prior state with the encoded future state. We perform this additional analysis to make sure that this effect does not only occur at the stimulus level, which is more directly linked to the objective of the predictive algorithm. This confirmatory analysis is shown in Figure 10, <u>Appendix B1.</u> (with tables in <u>Appendix C2.2.</u>). Since the actual PC operation subtracts the prior from the input, while our simple RNN models add the prior to the input, we expected the “prior” to be the negative latent state multiplied by the recurrent kernel. This transformation allows the PC subtraction to be implemented in a standard RNN architecture.

For statistical inference, we applied a Fisher-Z transformation to the correlation coefficients, inferred standard errors from the coefficients as described in [101], followed by Z-tests to compare all possible combinations of models with each other. We use Z-tests for this analysis to match the previously applied Fisher-Z transform. Confidence levels were corrected for all 36 possible between-model comparisons, using a Bonferroni correction.

#### 4.4.3. Learned representations

Any learning system should exhibit the capability to learn and form abstract representations [29,102,103]. Specifically, we are interested in whether the PC-inspired algorithms condense meaningful semantic information about the category of objects shown (in our case: numbers), without explicitly being trained to do so. This would demonstrate the network’s capacity to learn without supervision - a hallmark of biological neural learning systems such as PC [104–106].

To investigate learning performance of the networks, we examined the encoded information content in each of the models after being shown a single image. Each image was randomly augmented before feeding it to the network. Then, the encoded information was determined by using the latent output state of the network to decode the original number classes represented by the input. While the supervised condition has been explicitly trained to perform this task, for the predictive and the contrastive algorithms, no stimulus-class specific information has been fed to the network. Any better-than-untrained information in the network is therefore a result of the unsupervised learning procedures that these algorithms employ.

To encode information from the output state of the network, we used a simple logistic regression that classifies the original stimulus class based on the output features. From this, we calculated the overall decoding accuracy for all the stimulus classes for each model. For statistical inference, we calculated standard errors from the prediction accuracy using [107], followed by independent-sample T-tests. We performed these T-tests between all possible combinations of each of the 6 different models. Confidence levels were corrected for all 36 possible between-model comparisons, using a Bonferroni correction.

All the above tests were performed on the 6,000 test samples that were not used for training the models.

### 4.5. Analysis of Gain Control

We additionally investigated whether weight regularization can be used to simulate gain control. In PC, the noise level of the output signal is correlated with the noise level of the input. Gain control is the ability to manipulate the output noise level independently of the input noise [40,41]. Investigating gain control in our PC-inspired DNNs can provide insights into corresponding brain mechanisms. Weight regularization in neural networks is a widely known and used mechanic that constrains the magnitude of a network’s weights [108]. From preliminary investigations, we expected this to be a suitable mechanism to implement gain control, as weight regularization reduces the network’s sensitivity to variance in the input, potentially normalizing the statistical attributes of the output.

For this analysis, we trained all of the algorithms according to the standard setting without activity regularization, but with weight regularization. We compared the unregularized networks to the weight regularization networks. For weight regularized and unregularized networks of all algorithms, we passed single stimulus images while adding high (noise-to-signal standard deviation ratio ~1.7) or low (noise-to-signal standard deviation ratio ~0.3) Gaussian noise to the networks and collected the output activity. In order to create a distribution of variance ratios, we calculated the variances of the output activity and divided the variance of the high noise input images through the variances of the low noise input images for each sample. To evaluate whether the ratio of variance for both high and low noise images is significantly smaller for weight regularized networks than for unregularized networks (indicating gain control), we performed a Wilcoxon signed-rank test. We chose to use a nonparametric test for this analysis as the distribution of variance ratios was skewed and the requirement of normality was not given. Further we investigated whether despite weight regularization - there are still differences in variance distributions between low and high noise stimuli. We did this by evaluating if the variance distribution in the weight regularized conditions are indistinguishable between low noise and high noise stimuli. For this, we used another Wilcoxon signed-rank test. Significance levels were corrected for 6 multiple comparisons (2 tests with 3 comparisons) using a Bonferroni correction.

### 4.6. Other computational settings

Other details such as hyperparameters or software used are mentioned in Appendix A..

## Acknowledgments

We thank Mahdi Enan (Maastricht University) and Rasmus Bruckner (Hamburg University/Center for Cognitive Neuroscience Berlin) for feedback on this work. Further, we thank Johannes Singer (Center for Cognitive Neuroscience Berlin) for proofreading. This study was funded by the German Research Foundation / Deutsche Forschungsgemeinschaft (grant number AU 423/2-1).

## Appendix

### A. Technical details

#### A1. Network architecture

As mentioned in our methods (4. Methods), we used a typical RNN Network architecture for our model. Each of the RNN layers consisted of simple RNN (tf.keras.layers.SimpleRNN) combining an input kernel with a square recurrent kernel. The first layer in each network had 256 units, while the second had 128 units. All layers used leaky ReLU activation functions. The leaky ReLU activations were selected because they are bijective, enabling the encoded representations to be propagated back to the input level for facilitating the investigation of the network at a comprehensive level. Other than that, we used typical tensorflow default parameters, including glorot-uniform initializations for the feedforward kernel and orthogonal initializations for the recurrent kernel.

#### A2. Network and training hyperparameters

In addition to the different training procedures and the exploration of activity regularization, we also employed typical hyperparameter settings for the neural network models. As an optimization method, we used the Adam optimizer with the default TensorFlow 2.16 parameters. For the weight initializations, we employed Glorot-uniform initialization for the kernel weights and orthogonal initialization for the recurrent kernel weights. We use a batch size of 512 image sequences per batch. These fairly standard hyperparameter choices were made to ensure that any observed differences in the models’ performance and characteristics were primarily driven by the choice of training objective, rather than being confounded by other hyperparameter settings.

#### A3. Computational Environment

All network simulations in this study were performed using TensorFlow 2.12.0 [109], TensorFlow Addons 0.22 and TensorFlow Probability 0.20 and Keras 2.12.0 [110]. For the statistical analysis of the models’ performance and characteristics, we utilized scikit-learn 1.6.0 [111] and scipy 1.13.1 [112].

### B. Figures

#### B1. Prior correlation at encoding level

**Figure 10:**
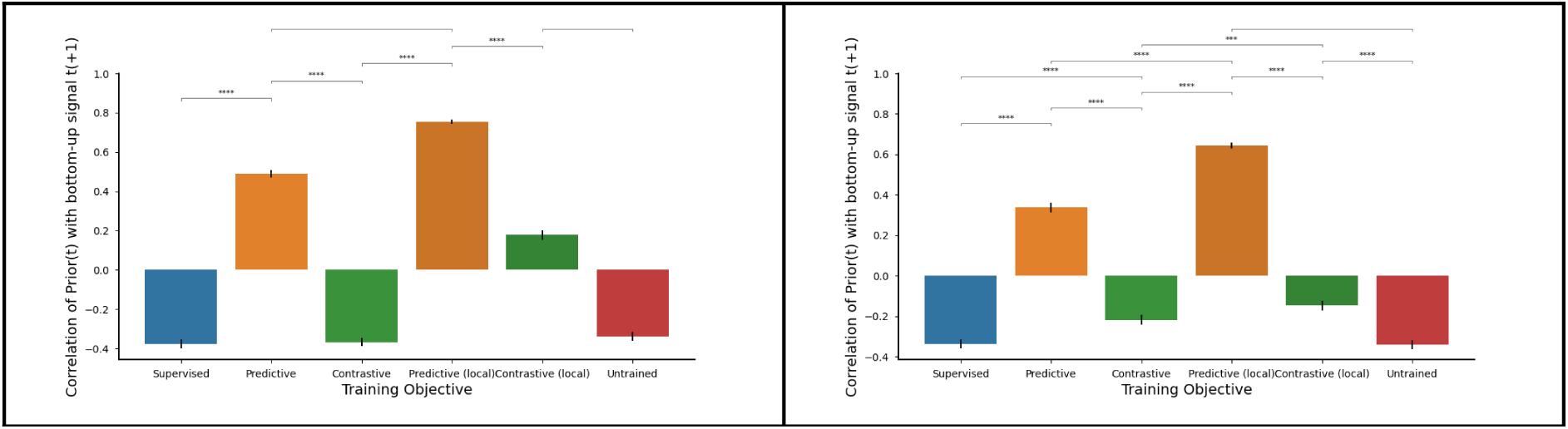
Correlations between priors and future input (encoded to the same level as priors). Left: No regularization, Right: With activity regularization.

#### B2. Predictive Condition MMR spread over layers

**Figure 11:**
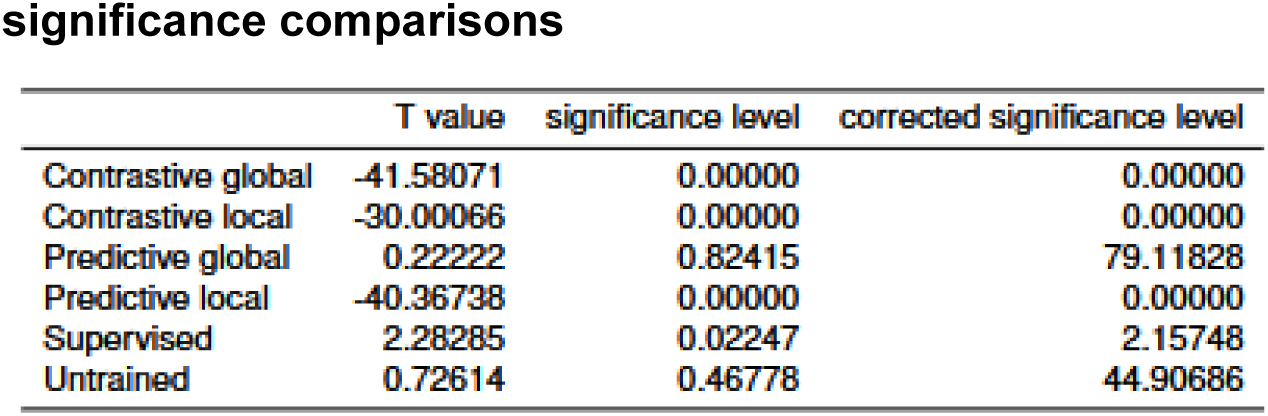
MMR for the “Predictive Global” condition, but separating the MMR for each specific layer. A. MMR for the first (lower) layer. B. MMR for the second (higher) layer.

### C. Tables

#### C1. Mismatch responses

##### No regularization

*Activity regularization* significance comparisons
|  | T value | significance level | corrected significance level |
| --- | --- | --- | --- |
| Contrastive global | -23.97432 | 0.00000 | 0.00000 |
| Contrastive local | -15.89864 | 0.00000 | 0.00000 |
| Predictive global | -11.68170 | 0.00000 | 0.00000 |
| Predictive local | -20.63438 | 0.00000 | 0.00000 |
| Supervised | 17.60957 | 0.00000 | 0.00000 |
| Untrained | 0.72614 | 0.46778 | 44.90686 |

##### Activity regularization

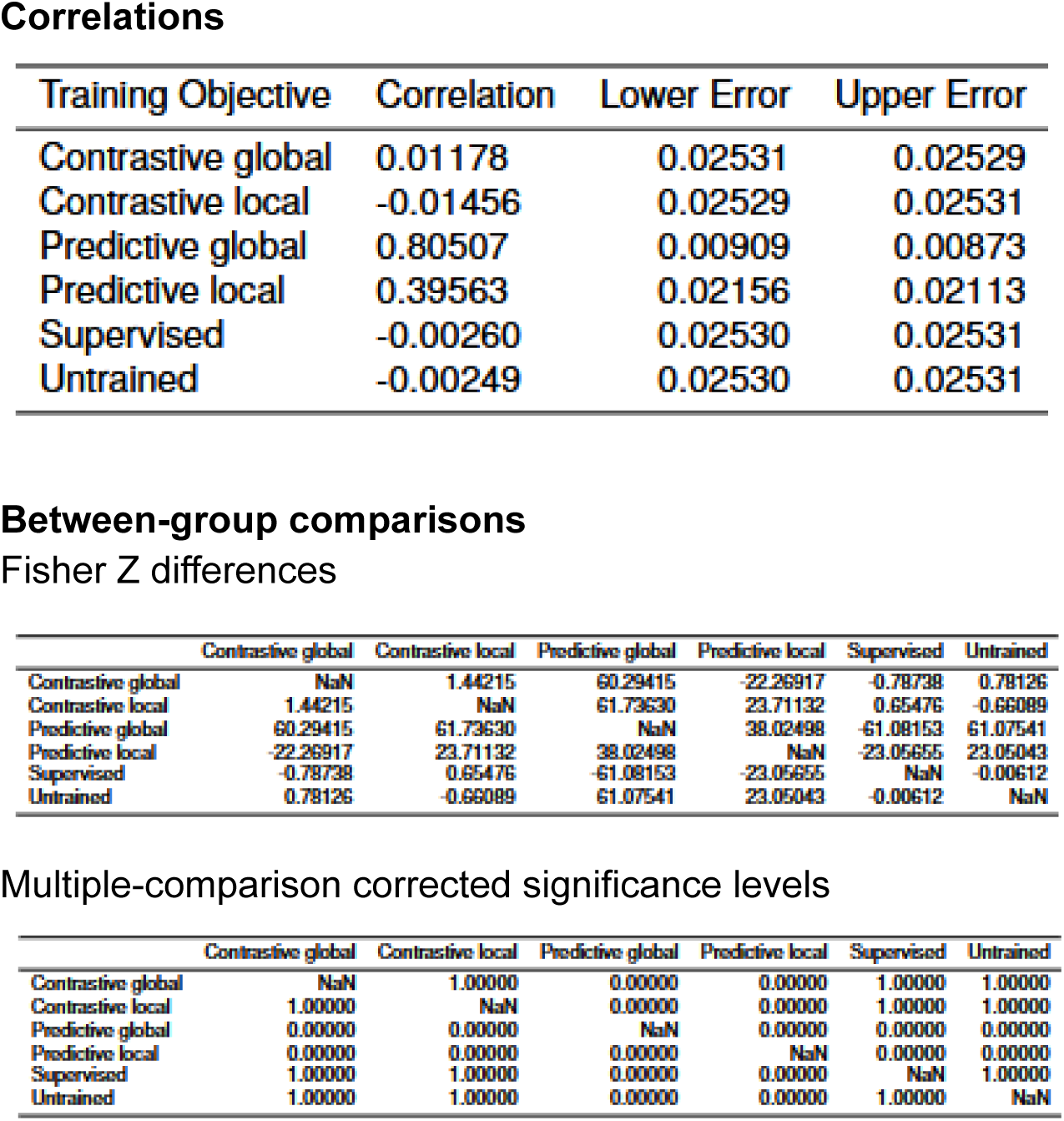

Correlations
| Training Objective | Correlation | Lower Error | Upper Error |
| --- | --- | --- | --- |
| Contrastive global | 0.01178 | 0.02531 | 0.02529 |
| Contrastive local | -0.01456 | 0.02529 | 0.02531 |
| Predictive global | 0.80507 | 0.00909 | 0.00873 |
| Predictive local | 0.39563 | 0.02156 | 0.02113 |
| Supervised | -0.00260 | 0.02530 | 0.02531 |
| Untrained | -0.00249 | 0.02530 | 0.02531 |

Between-group comparisons Fisher Z differences
|  | Contrastive global | Contrastive local | Predictive global | Predictive local | Supervised | Untrained |
| --- | --- | --- | --- | --- | --- | --- |
| Contrastive global | NaN | 1.44215 | 60.29415 | -22.26917 | -0.78738 | 0.78126 |
| Contrastive local | 1.44215 | NaN | 61.73630 | 23.71132 | 0.65476 | -0.66089 |
| Predictive global | 60.29415 | 61.73630 | NaN | 38.02498 | -61.08153 | 61.07541 |
| Predictive local | -22.26917 | 23.71132 | 38.02498 | NaN | -23.05655 | 23.05043 |
| Supervised | -0.78738 | 0.65476 | -61.08153 | -23.05655 | NaN | -0.00612 |
| Untrained | 0.78126 | -0.66089 | 61.07541 | 23.05043 | -0.00612 | NaN |

Multiple-comparison corrected significance levels
|  | Contrastive global | Contrastive local | Predictive global | Predictive local | Supervised | Untrained |
| --- | --- | --- | --- | --- | --- | --- |
| Contrastive global | NaN | 1.00000 | 0.00000 | 0.00000 | 1.00000 | 1.00000 |
| Contrastive local | 1.00000 | NaN | 0.00000 | 0.00000 | 1.00000 | 1.00000 |
| Predictive global | 0.00000 | 0.00000 | NaN | 0.00000 | 0.00000 | 0.00000 |
| Predictive local | 0.00000 | 0.00000 | 0.00000 | NaN | 0.00000 | 0.00000 |
| Supervised | 1.00000 | 1.00000 | 0.00000 | 0.00000 | NaN | 1.00000 |
| Untrained | 1.00000 | 1.00000 | 0.00000 | 0.00000 | 1.00000 | NaN |

#### C2. Prior expectations

##### C2.1. Input level

###### No regularization

Activity regularization Correlations
| Training Objective | Correlation | Lower Error | Upper Error |
| --- | --- | --- | --- |
| Predictive global | 0.78028 | 0.01010 | 0.00971 |
| Contrastive local | -0.00059 | 0.02530 | 0.02530 |
| Contrastive global | 0.00826 | 0.02531 | 0.02530 |
| Predictive local | 0.57463 | 0.01720 | 0.01671 |
| Supervised | -0.01706 | 0.02529 | 0.02531 |
| Untrained | -0.00249 | 0.02530 | 0.02531 |

Between-group comparisons Fisher Z differences
|  | Contrastive global | Contrastive local | Predictive global | Predictive local | Supervised | Untrained |
| --- | --- | --- | --- | --- | --- | --- |
| Contrastive global | NaN | 0.48469 | 56.82979 | -35.38229 | -1.38661 | 0.58876 |
| Contrastive local | 0.48469 | NaN | 57.31448 | 35.86698 | -0.90192 | 0.10407 |
| Predictive global | 56.82979 | 57.31448 | NaN | 21.44750 | -58.21640 | 57.41855 |
| Predictive local | -35.38229 | 35.86698 | 21.44750 | NaN | -36.76890 | 35.97105 |
| Supervised | -1.38661 | -0.90192 | -58.21640 | -36.76890 | NaN | -0.79785 |
| Untrained | 0.58876 | 0.10407 | 57.41855 | 35.97105 | -0.79785 | NaN |

Multiple-comparison corrected significance levels

###### Activity regularization

Correlations
| Training Objective | Correlation | Lower Error | Upper Error |
| --- | --- | --- | --- |
| Contrastive global | -0.36864 | 0.02166 | 0.02207 |
| Contrastive local | 0.17834 | 0.02461 | 0.02439 |
| Predictive global | 0.48990 | 0.01947 | 0.01900 |
| Predictive local | 0.75453 | 0.01111 | 0.01069 |
| Supervised | -0.37684 | 0.02151 | 0.02192 |
| Untrained | -0.34057 | 0.02218 | 0.02256 |

Between-group comparisons Fisher Z differences
|  | Contrastive global | Contrastive local | Predictive global | Predictive local | Supervised | Untrained |
| --- | --- | --- | --- | --- | --- | --- |
| Contrastive global | NaN | -31.05411 | 50.52997 | -75.03205 | -0.52167 | -1.75800 |
| Contrastive local | -31.05411 | NaN | 19.47586 | 43.97794 | -31.57578 | 29.29612 |
| Predictive global | 50.52997 | 19.47586 | NaN | -24.50208 | -51.05164 | 48.77198 |
| Predictive local | -75.03205 | 43.97794 | -24.50208 | NaN | -75.55372 | 73.27406 |
| Supervised | -0.52167 | -31.57578 | -51.05164 | -75.55372 | NaN | -2.27967 |
| Untrained | -1.75800 | 29.29612 | 48.77198 | 73.27406 | -2.27967 | NaN |

Multiple-comparison corrected significance levels
|  | Contrastive global | Contrastive local | Predictive global | Predictive local | Supervised | Untrained |
| --- | --- | --- | --- | --- | --- | --- |
| Contrastive global | NaN | 0.00000 | 0.00000 | 0.00000 | 1.00000 | 1.00000 |
| Contrastive local | 0.00000 | NaN | 0.00000 | 0.00000 | 0.00000 | 0.00000 |
| Predictive global | 0.00000 | 0.00000 | NaN | 0.00000 | 0.00000 | 0.00000 |
| Predictive local | 0.00000 | 0.00000 | 0.00000 | NaN | 0.00000 | 0.00000 |
| Supervised | 1.00000 | 0.00000 | 0.00000 | 0.00000 | NaN | 0.33941 |
| Untrained | 1.00000 | 0.00000 | 0.00000 | 0.00000 | 0.33941 | NaN |

##### C2.2. Encoded level

###### No regularization

Activity regularization Correlations
| Training Objective | Correlation | Lower Error | Upper Error |
| --- | --- | --- | --- |
| Contrastive global | -0.21764 | 0.02397 | 0.02424 |
| Contrastive local | -0.14674 | 0.02467 | 0.02485 |
| Predictive global | 0.33751 | 0.02261 | 0.02223 |
| Predictive local | 0.64315 | 0.01508 | 0.01460 |
| Supervised | -0.33657 | 0.02225 | 0.02263 |
| Untrained | -0.34057 | 0.02218 | 0.02256 |

Between-group comparisons Fisher Z differences
|  | Contrastive global | Contrastive local | Predictive global | Predictive local | Supervised | Untrained |
| --- | --- | --- | --- | --- | --- | --- |
| Contrastive global | NaN | -4.01783 | 31.34730 | -53.92155 | -7.06582 | 7.31347 |
| Contrastive local | -4.01783 | NaN | 27.32947 | 49.90372 | -11.08365 | 11.33130 |
| Predictive global | 31.34730 | 27.32947 | NaN | -22.57425 | -38.41311 | 38.66076 |
| Predictive local | -53.92155 | 49.90372 | -22.57425 | NaN | -60.98737 | 61.23502 |
| Supervised | -7.06582 | -11.08365 | -38.41311 | -60.98737 | NaN | 0.24765 |
| Untrained | 7.31347 | 11.33130 | 38.66076 | 61.23502 | 0.24765 | NaN |

Multiple-comparison corrected significance levels
|  | Contrastive global | Contrastive local | Predictive global | Predictive local | Supervised | Untrained |
| --- | --- | --- | --- | --- | --- | --- |
| Contrastive global | NaN | 0.00088 | 0.00000 | 0.00000 | 0.00000 | 0.00000 |
| Contrastive local | 0.00088 | NaN | 0.00000 | 0.00000 | 0.00000 | 0.00000 |
| Predictive global | 0.00000 | 0.00000 | NaN | 0.00000 | 0.00000 | 0.00000 |
| Predictive local | 0.00000 | 0.00000 | 0.00000 | NaN | 0.00000 | 0.00000 |
| Supervised | 0.00000 | 0.00000 | 0.00000 | 0.00000 | NaN | 1.00000 |
| Untrained | 0.00000 | 0.00000 | 0.00000 | 0.00000 | 1.00000 | NaN |

###### Activity regularization

Accuracies
| Training Objective | Accuracy | Lower Error | Upper Error |
| --- | --- | --- | --- |
| Contrastive global | 0.36550 | 0.01219 | 0.01219 |
| Contrastive local | 0.35217 | 0.01209 | 0.01209 |
| Predictive global | 0.38667 | 0.01232 | 0.01232 |
| Predictive local | 0.34700 | 0.01204 | 0.01204 |
| Supervised | 0.36667 | 0.01219 | 0.01219 |
| Untrained | 0.32067 | 0.01181 | 0.01181 |

Between-group comparisons T values
|  | Contrastive global | Contrastive local | Predictive global | Predictive local | Supervised | Untrained |
| --- | --- | --- | --- | --- | --- | --- |
| Contrastive global | NaN | 1.52268 | 2.39393 | 2.11630 | 0.13265 | 5.17835 |
| Contrastive local | 1.52268 | NaN | 3.91767 | -0.59348 | 1.65536 | 3.65364 |
| Predictive global | 2.39393 | 3.91767 | NaN | 4.51192 | -2.26123 | 7.57907 |
| Predictive local | 2.11630 | -0.59348 | 4.51192 | NaN | 2.24899 | 3.05970 |
| Supervised | 0.13265 | 1.65536 | -2.26123 | 2.24899 | NaN | 5.31126 |
| Untrained | 5.17835 | 3.65364 | 7.57907 | 3.05970 | 5.31126 | NaN |

Multiple-comparison corrected significance levels
|  | Contrastive global | Contrastive local | Predictive global | Predictive local | Supervised | Untrained |
| --- | --- | --- | --- | --- | --- | --- |
| Contrastive global | NaN | 1.00000 | 0.25026 | 0.51510 | 1.00000 | 0.00000 |
| Contrastive local | 1.00000 | NaN | 0.00135 | 1.00000 | 1.00000 | 0.00389 |
| Predictive global | 0.25026 | 0.00135 | NaN | 0.00010 | 0.35644 | 0.00000 |
| Predictive local | 0.51510 | 1.00000 | 0.00010 | NaN | 0.36796 | 0.03331 |
| Supervised | 1.00000 | 1.00000 | 0.35644 | 0.36796 | NaN | 0.00000 |
| Untrained | 0.00000 | 0.00389 | 0.00000 | 0.03331 | 0.00000 | NaN |

##### C3. Learned representations

###### No regularization

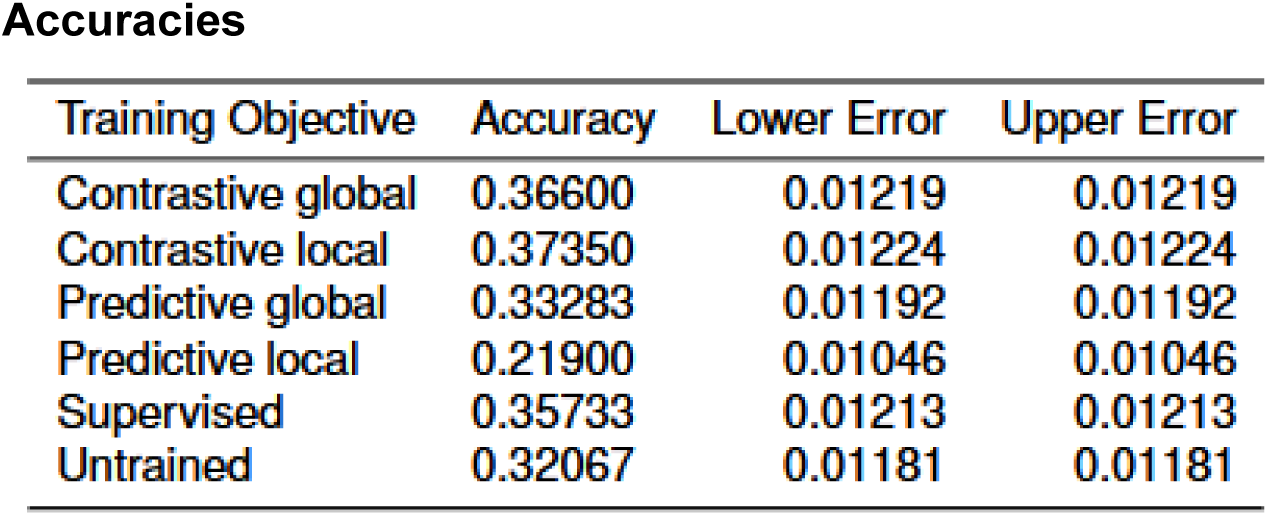

*Activity regularization* Accuracies
| Training Objective | Accuracy | Lower Error | Upper Error |
| --- | --- | --- | --- |
| Contrastive global | 0.36600 | 0.01219 | 0.01219 |
| Contrastive local | 0.37350 | 0.01224 | 0.01224 |
| Predictive global | 0.33283 | 0.01192 | 0.01192 |
| Predictive local | 0.21900 | 0.01046 | 0.01046 |
| Supervised | 0.35733 | 0.01213 | 0.01213 |
| Untrained | 0.32067 | 0.01181 | 0.01181 |

###### Activity regularization

Between-group comparisons T values
|  | Contrastive global | Contrastive local | Predictive global | Predictive local | Supervised | Untrained |
| --- | --- | --- | --- | --- | --- | --- |
| Contrastive global | NaN | -0.85099 | -3.81243 | 17.93477 | -0.98799 | 5.23532 |
| Contrastive local | -0.85099 | NaN | -4.66452 | -18.80429 | -1.83909 | 6.08825 |
| Predictive global | -3.81243 | -4.66452 | NaN | 14.06361 | 2.82368 | 1.42093 |
| Predictive local | 17.93477 | -18.80429 | 14.06361 | NaN | 16.92781 | -12.62837 |
| Supervised | -0.98799 | -1.83909 | 2.82368 | 16.92781 | NaN | 4.24578 |
| Untrained | 5.23532 | 6.08825 | 1.42093 | -12.62837 | 4.24578 | NaN |

Multiple-comparison corrected significance levels
|  | Contrastive global | Contrastive local | Predictive global | Predictive local | Supervised | Untrained |
| --- | --- | --- | --- | --- | --- | --- |
| Contrastive global | NaN | 1.00000 | 0.00207 | 0.00000 | 1.00000 | 0.00000 |
| Contrastive local | 1.00000 | NaN | 0.00005 | 0.00000 | 0.98889 | 0.00000 |
| Predictive global | 0.00207 | 0.00005 | NaN | 0.00000 | 0.07133 | 1.00000 |
| Predictive local | 0.00000 | 0.00000 | 0.00000 | NaN | 0.00000 | 0.00000 |
| Supervised | 1.00000 | 0.98889 | 0.07133 | 0.00000 | NaN | 0.00033 |
| Untrained | 0.00000 | 0.00000 | 1.00000 | 0.00000 | 0.00033 | NaN |

##### C4. Gain control

| Algorithm | Variance ratio W | Variance ratio Z | Variance ratio P | Variance ratio P (corrected) | Regularization W | Regularization Z | Regularization P | Regularization P (corrected) |
| --- | --- | --- | --- | --- | --- | --- | --- | --- |
| 0 Contrastive | 0 | -67.08483 | 0.00000 | 0.00000 | 7 | -67.08478 | 0.00000 | 0.00000 |
| 1 Predictive | 323061 | -64.67718 | 0.00000 | 0.00000 | 20846 | -66.92948 | 0.00000 | 0.00000 |
| 2 Supervised | 8 | -67.08477 | 0.00000 | 0.00000 | 4144198 | -36.19967 | 0.00000 | 0.00000 |

